# Acute aggressive behavior perturbates the oxidative status of a wild bird independently of testosterone and progesterone

**DOI:** 10.1101/2020.04.06.027029

**Authors:** Lucia Mentesana, Nicolas M. Adreani

## Abstract

Aerobically demanding activities like aggression can lead to an elevated oxidative metabolism affecting the concentration of pro-oxidant and antioxidant compounds and can result in an overall perturbation of the oxidative status. Aggression may also alter the oxidative status indirectly through an increase in testosterone and progesterone concentrations. Given that changes in the oxidative status could represent a physiological cost of aggression, we tested the hypothesis that acute conspecific aggression impairs the oxidative status and evaluated the role of testosterone and progesterone as potential mediators. To achieve this, we experimentally manipulated the aggressive behavior of wild female and male birds and measured the concentrations of pro-oxidants, enzymatic- and non-enzymatic antioxidants, testosterone and progesterone in blood. After 20 minutes of conspecific aggressive behavior, both sexes had lower concentrations of non-enzymatic antioxidants than control individuals. This effect was independent of testosterone and progesterone concentrations, and much stronger in females than in males. Further, only in females (but not in males) being more aggressive came at the expense of lower antioxidant concentration. We provide the first experimental evidence that acute aggressive behavior perturbates the oxidative state of a wild vertebrate independently of testosterone and progesterone, with potential ecological and evolutionary implications given the role of the redox system in shaping life-history traits.

## 1. Introduction

Animals frequently engage in aggressive behaviors to obtain and secure limited resources that can enhance their reproductive success, such as a breeding territory (Beckman & Ames, 1998; Duckworth, 2006; Demas et al. 2007; Rosvall, 2008; Smith & Blumstein 2008; Clutton-Brock, 2009). However, aggressive behaviors entail costs (e.g., territory loss or injuries), which could partially explain why individuals differ in their degree of aggressiveness. A particularly relevant but understudied cost associated with this behavior concerns the oxidative state of an individual (Costantini et al., 2008; Isaksson et al., 2011). The oxidative state of an individual is determined by the concentration of pro-oxidants (i.e., reactive oxygen species) and antioxidants (i.e., non-enzymatic and enzymatic compounds) present in cells and tissues (reviewed by Costantini 2019). A change in any of these molecular components in favor of pro-oxidants can lead to damage of biomolecules such as lipids, proteins and DNA (reviewed by Costantini 2008; Monaghan et al., 2009). This damage can shape life-history decisions of individuals as well as life history traits such as reproduction and longevity (e.g., Finkel and Holbrook, 2000; Costantini 2008; Monaghan et al. 2009), potentially translating into ecological and evolutionary consequences.

Aggressive behaviors are energetically demanding activities that increase the metabolic rate, exposing an individual to an elevated concentration of pro-oxidants (Figure 1a; Costantini 2008; Powers & Jackson, 2008; Skrip & McWilliams, 2016; Cooper-Mullin & McWilliams, 2016; but see Salin et al., 2015). Such perturbations in the oxidative status of animals are expected since more than 90% of the cellular energy is generated by the mitochondria (Bottje 2015) and natural by-products of aerobic respiration are reactive oxygen species (ROS). However, our understanding of aggression altering the oxidative state is still in its early stages. In selected lines of mice (*Mus musculus domesticus*), aggressive males had lower levels of antioxidants than non-aggressive males (Costantini et al., 2008). In contrast, aggressive males of wild-caught white skinks (*Egernia whitii*) had higher levels of non-enzymatic antioxidants than less aggressive males, whereas in females no such relationship was apparent (Isaksson et al., 2011). Acute aggressive behaviors, such as territorial fights, that represent an increase in energy expenditure, could also incur an oxidative challenge; yet it remains unknown whether this is the case.

**Figure 1.**
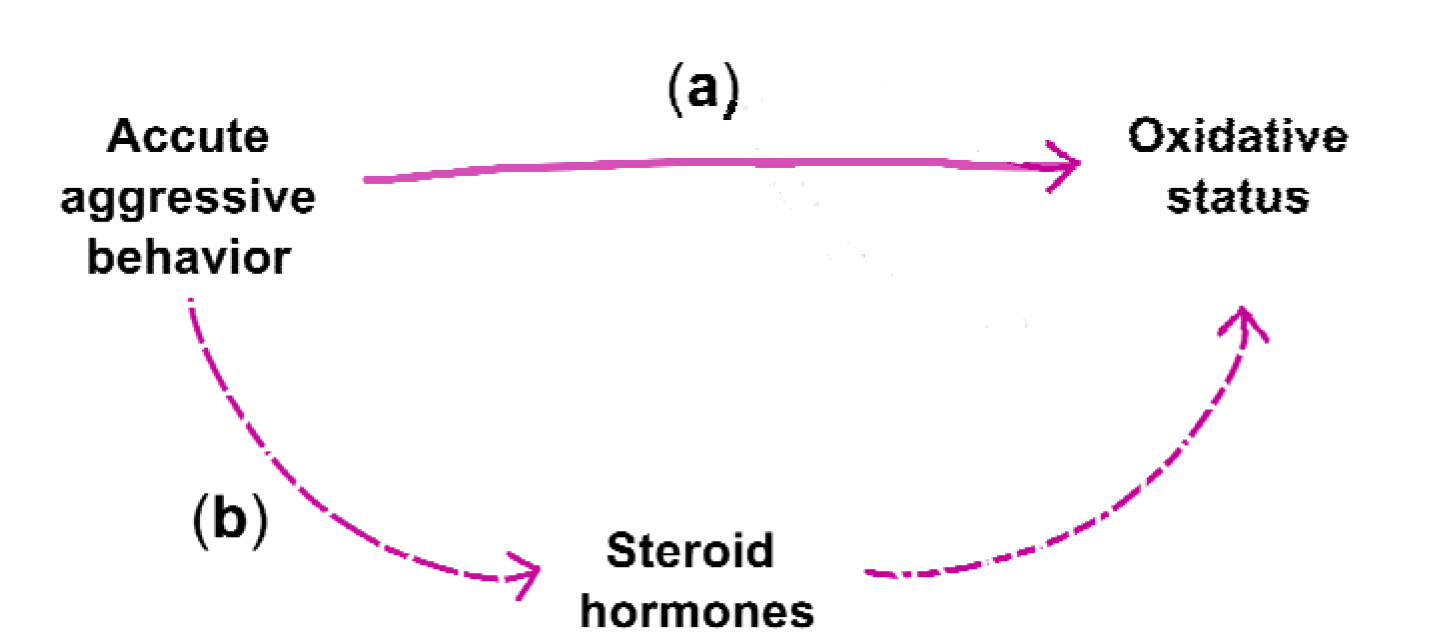
Hypothesized pathways by which acute aggressive behaviors can perturbate the individuals’ oxidative status: (a) directly through an increase in the metabolic rate, and/or (b) indirectly through the action of steroid hormones.

The oxidative state of an individual can be indirectly altered by aggressive behaviors through an increase in steroid hormone concentrations (Figure 1b; Schantz et al., 1999; Alonso-Alvarez et al., 2007). The sex steroid testosterone is assumed to be the key hormone related to resource-defense aggression and can increase during aggressive interactions in males (Wingfield et al. 1990, Hirchenhauser et al., 2006; Goymann et al., 2007; Hau 2007). Testosterone has been proposed to be a cause of increased ROS production (e.g., Alonso-Alvarez et al., 2007, Koch et al. 2016), which can occur either because testosterone enhances the metabolic rate of individuals (e.g., Marler & Moore, 1989; Welle et al. 1992; Wikelsky et al., 1999; Buchanan et al. 2001, Koch et al. 2016) and/or because testosterone has intrinsic oxidative properties (e.g., Zhu et al., 1997; Chainy et al., 2009; Casagrande et al. 2011). An increase in aggressive behavior after a territorial intrusion can, therefore, potentially lead to an increase in testosterone concentrations (e.g., Wingfield & Wada 1989; Wingfield & Hahn, 1994; McGlothlin et al., 2008), with testosterone levels influencing the concentrations of pro-oxidant and antioxidant compounds and resulting in an overall perturbation of the oxidative state. To date, only one study has looked at the relationship between aggression, oxidative status and testosterone concentrations in a wild-caught vertebrate (Isaksson et al., 2011). Isaksson et al (2011) found that the aggressive phenotype of male white skinks, but not testosterone concentrations, was positively related to the oxidative condition of individuals. Aggression can also be associated with another steroid hormone: progesterone. Progesterone has been mainly proposed as a mediator of aggressive behavior in females (e.g., Goymann et al., 2008; but see e.g., Adreani et al., 2018). As with testosterone, progesterone can increase the metabolic rate of organisms (Gavrilova-Jordan & Price 2007) and the concentration of pro-oxidants (only from studies done *in vitro* e.g., Zhu et al., 1997; Itagaki et al., 2005). Investigating the relationships between aggressive behavior, oxidative status, testosterone and progesterone in females and males is fundamental if we aim to understand the sex-specificity of these physiological pathways.

The main goal of our study was to test the hypothesis that an acute increase in conspecific aggressive behavior can impair the oxidative status of wild rufous horneros (Aves: *Furnarius rufus*, hereafter termed hornero), and to test whether this was related to testosterone and progesterone concentrations. To elicit an aggressive response, we challenged female and male birds with 20 minutes of simulated conspecific territorial intrusions (STI) during the nest-building period. Immediately after the STI we collected blood samples to measure the concentrations of three oxidative status markers, testosterone and progesterone. We then compared these concentrations with the ones of birds that did not engage in aggressive behaviors. The hornero represents an excellent model system to study the effects of acute aggression across sexes. It is a seasonal breeder, and both females and males are involved in territorial defense throughout the year (Fraga, 1980; Diniz et al., 2016; Mentesana et al. 2020). Further, horneros are sexually monomorphic in plumage coloration and body size (i.e., there is no difference in body condition between sexes; Diniz et al., 2016), and all breeding behaviors studied so far, except for aggressive behaviors, are shared and coordinated between the sexes (Massoni et al., 2012; Diniz et al. 2018; Mentesana et al. 2020).

## 2. Materials and methods

### 2.1 Field site and simulated territorial intrusion (STI)

The study was conducted between August 22nd and September 28th of 2016 on the campus of INIA ‘Las Brujas’ (National Institute of Agricultural Research), Department of Canelones, Uruguay (34°40’ S, 56°20’ W; 0-35 m a.s.l.). During this period, horneros were building their nests and females were close to egg laying. In total, we collected behavioral and physiological data from birds defending 51 territories that were subjected to the experimental treatments between 7 am and 1 pm.

We experimentally manipulated the hornero’s aggressive behavior by performing simulated territorial intrusions (STI). For each territory, the treatment (i.e., control or STI) was randomly determined. STI is a common method to investigate the aggressive and physiological responses of a territory holder in response to an intruder (e.g., Gill et al., 2007; Apfelbeck & Goymann, 2011; Villavicencio et al., 2014), and STIs are known to be effective in horneros (Adreani et al., 2018; Mentesana et al., 2020). In particular, a stuffed decoy of a male hornero, a speaker and folded mist nets were placed 5 m away from the nest of focal pairs, and solo songs and duets of conspecifics were played (recordings were extracted from the database www.xeno-canto.org and the amplitude of every recording was normalized). For ethical reasons we could not use more than one decoy (i.e., the visual component of the STIs was the same for every territory). However, horneros are sexually indistinguishable in morphological traits (i.e., mass, tarsus and bill) and plumage coloration (assessed in relation to the avian visual system; Diniz et al., 2016). Further, in a comparable STI study done in European great tits (*Parus major*) the decoys used (n=15) did not explain any significant variation in aggressive behaviors (Araya-Ajoy et al., 2013); thus, suggesting that the use of only one decoy did not bias our results. On the other hand, given the lack of information on the function of the different vocal signals in horneros, we decided to form a pool of ten male solo songs and ten duets to ensure that both females and males responded to the STI. We also added ten ‘silence’ files of 7-15 sec duration to have unpredictable gaps between the different stimuli. The auditory stimuli for each STI were randomly selected from this pool to avoid pseudo-replication of the acoustic component across territories (e.g., Apfelbek et al., 2011). Horneros are suboscines and do not learn their vocalizations (Freeman et al., 2017); compared to oscines, the acoustic variability of songs and duets across individuals is low (Freeman et al., 2017). Hence, it is likely that all playbacks elicited a comparable behavioral response across territories despite the auditory stimuli being randomly chosen from the pool.

Challenged birds (hereafter ‘STI birds’, N_females_, N_males_ = 9, 18) were exposed to a simulated territorial intrusion by a conspecific for 20 minutes. We decided on this experimental time because, in birds, plasma levels of testosterone increase after 10 minutes of the onset of the challenge and can remain high until an hour after onset of the intrusion (Wingfield & Wada 1989). During the 20 minutes of territorial intrusion, two observers recorded the aggressive behavior of the territorial couple in digital voice recorders (Philips VoiceTraicer DVT1200 and Olympus Digital Recorder VN-733 PC). Each observer followed one member of the focal STI pair. Which individual of the pair to follow was randomly determined prior to the experimental trial. Sex identification in the field was done based on the singing behavior (i.e., the contribution of females and males in the duets are acoustically distinct from each other; Laje & Mindlin 2003) and was later confirmed by molecular sexing (for details see Adreani et al., 2018). Observers did not differ in their quantification skills for the behaviors scored (Figure S1; Table S1; see Supplementary Material section for a detailed description of the inter-observer reliability test). The following parameters were recorded: 1) response latency (time between start of playback and first approach), 2) time spent within 5 m of the decoy, 3) time spent on the nest, 4) number of solo songs, 5) number of duets, and 6) number of flights over the decoy (which represent both attacks and attempted attacks towards the decoy). After 20 minutes, mist nets were unfolded for a maximum of ten additional minutes. Capturing horneros passively is of great difficulty (LM & NMA, personal observations). Therefore, to attract control individuals (N_females_, N_males_ = 7, 24) we used a stuffed decoy of a male hornero and playbacks, but mist nets were kept open since the beginning of the stimuli presentation. That is, control birds did not perform aggressive behaviors since they were caught as soon as they approached the experimental set-up. Control birds were captured within the first minutes of exposure to the decoy and playback (mean capture time ± sd = 4.57 ± 2.97 mins for females and 4.50 ± 3.75 mins for males), while STI birds were captured after 20 minutes of exposure (mean capture time ± sd = 23.11 ± 2.33 mins for females and 25.94 ± 4.68 mins for males; Figure S2). From all captured birds, 80 μl of blood were taken from the brachial vein using heparinized capillaries. In addition, morphometric measurements (i.e. mass, tarsus length and wing length) were recorded. Control and STI individuals did not differ in body condition index (Table S2). Before releasing them back in their territories, each individual was marked with a numbered aluminum ring and a unique combination of three colored plastic split rings.

Horneros respond immediately towards the acoustic and visual stimuli and approach the decoy (Adreani et al., 2018; Mentesana et al., 2020). In our experiments STIs elicited similar behavioral responses as intrusions by real conspecifics or territorial conflicts with neighbours (LM & NMA, personal observations). Females and males showed clear behavioral responses towards our treatment. However, once the male was captured, the female would immediately stop any territorial behavior and start alarm calling. This explains why, during our experiments we caught more than twice the number of males compared to females, and why we generally caught only one member of the pair.

### 2.2 Oxidative status analyses

The oxidative status of female and male horneros was determined from both non-enzymatic and enzymatic antioxidants present in plasma and red blood cells respectively, and from reactive oxygen species present in plasma. Non-enzymatic antioxidant concentrations (OXY) were measured using the OXY-Adsorbent test (Diacron International SRL, Grosseto, Italy; Costantini et al. (2006). This assay quantifies the ability of antioxidants to cope with the oxidant action of hypochlorous acid. Enzymatic antioxidant concentrations of glutathione peroxidase (GPX) were determined using the Ransel assay (Randox Laboratories, Germany; Costantini et al. 2011). This assay determines the activity of the enzyme when catalyzing the oxidation of glutathione by cumune hydroperoxide. Oxidative damage was quantified by measuring the concentrations of reactive oxygen metabolites (ROMs) with the d-ROM test (Diacron International SRL, Grosseto, Italy; Costantini et al. 2006). ROMs are end-products of free radical reactions with biological macromolecules. The absorbance was read using a spectrophotometer (Thermo Scientific Multiskan GO) at a wavelength of 546 nm for OXY and ROMs (endpoint mode) and 340 nm for GPX (kinetic modality).

### 2.3 Testosterone and progesterone analyses

Testosterone and progesterone hormones were extracted from plasma, and concentrations were determined by radioimmunoassay following Goymann et al. (2008). Females and males were analyzed in two separate assays because the samples collected for males were also part of a complementary study (for details see Adreani et al. 2018). In females, mean recoveries following extractions of the samples were (mean ± sd) 89 ± 2% and 80 ± 3% for testosterone and progesterone, respectively. The lower detection limits of the testosterone and progesterone assay were 0.39 and 0.38 pg/tube. For both hormones, all samples used in our analyses were above these detection limits. The intra-assay variation, calculated from of an extracted chicken pool, were 7.5% for testosterone and 12.2% for progesterone. It is worth noting that, despite analyzing female and male samples in separate assays, mean recoveries for both hormones were similar between sexes (male mean recovery of testosterone ± sd = 87 ± 1.9 % and progesterone = 84 ± 5). It is therefore unlikely that hormonal differences between females and males are due to samples measured in separate assays.

### 2.4 Statistical analysis

All our analyses were performed using the R-packages “lme4” and “arm” in R-3.3.3 (R Core Team, 2013) in a Bayesian framework with non-informative priors. For all models, we assumed a Gaussian error distribution, and model fit was assessed by visual inspection of the residuals. We used the “*sim*” function to simulate posterior distributions of the model parameters. Based on 10,000 simulations, we extracted the mean value and 95% credible intervals (CrI) (Gelman & Hill, 2007). Assessment of statistical support was obtained from the posterior distribution of each parameter (Korner-Nievergelt et al., 2015). We considered an effect to be statistically meaningful when the posterior probability of the mean difference (hereafter termed ‘*p*(*dif*)’) between compared estimates was higher than 95% or when the estimated CrI did not include zero. For details on this approach see Korner-Nievergelt et al. (2015).

First, we ran five linear models fitting OXY, GPX, ROMs, testosterone and progesterone as response variables to i) compare the oxidative status, testosterone and progesterone concentrations between female and male control birds (i.e., individuals that better represent the baseline status), and to ii) study the effect of 20 minutes of aggressive behavior on the oxidative status, testosterone and progesterone concentrations of female and male horneros. To do so, sex (female vs male) and treatment (control vs STI) were included as fixed effects, as well as the interaction between both factors. We also included capture time and handling time as covariates to account for the possible variation that they might add (see Table S3). Preliminary exploration of our data showed that time of the day as well as observer ID did not explain significant variance; thus, to avoid overparameterization, they were not included in the final model.

Second, we ran three linear models to investigate the relationship between oxidative status and testosterone concentrations in control individuals and three similar models to investigate the relationship between oxidative status and progesterone. In the first set of models OXY, GPX or ROMs were fitted as response variables and testosterone as explanatory variable. In the second set of models, progesterone was fitted as explanatory variable. In all models, sex and the interaction of sex with the hormonal levels were included as fixed factors. Since females and males differ in their testosterone concentrations (see Figure 1), for each sex we normalized the hormonal levels between 0 and 1. This allowed us to compare results between sexes.

Finally, within the STI birds, we studied the effect that aggressive behavior had on the oxidative status, testosterone and progesterone concentrations. To do so, we fitted OXY, GPX, ROMs, testosterone and progesterone concentrations as the response variable in separate linear models. While GPX and ROMs were square-root transformed for a better fit of each model, testosterone was log-transformed for the same purpose. Although we measured 6 behavioral responses as proxies of aggressiveness (detailed above), to avoid multiple testing, here we only used “flights over the decoy” as explanatory variable. We followed this approach based on a recent study that shows that in horneros the 6 behavioral responses are to some extent correlated, but “flights over the decoy” is the best proxy for conspecific territorial aggression during the breeding season (Mentesana et al. 2020). Male horneros are more aggressive than females (Diniz et al. 2018; Mentesana et al. 2020); thus, for each sex, we normalized the behavioral variables between 0 and 1 in order to capture the aggression gradient in each sex and to be able to compare the effects between the sexes in one model. Sex and the interaction of this variable with aggressive behavior were fitted as fixed factors. Given that we had a limited sample size and that neither capture nor handling time had an effect on any of the oxidative parameters or testosterone levels (see Table S3), these two variables were not included in the final models.

## 3. Results

### 3.1 Sex differences in the concentrations of oxidative markers, testosterone and progesterone of control birds

Control females had a lower non-enzymatic antioxidant capacity (OXY) than control males (Figure 2A; *p*(*dif*.) > 99.99%, Table S3). Both sexes had similar enzymatic antioxidant concentrations (GPX; Figure 2B; *p*(*dif*.) = 79.03%, Table S3). We found a trend for control males having lower levels of pro-oxidants than females (ROMs; Figure 2C; *p*(*dif*.) = 94.67%, Table S3). Control females had lower testosterone concentrations than control males (Figure 2D; *p*(*dif*.) = 99.86%, Table S3), but we found no difference in progesterone concentrations between sexes (Figure 2E; *p*(*dif*.) = 62.51%, Table S3). There was no relationship between any oxidative marker and testosterone or progesterone concentrations in either sex (Figure 3, Table S4 Table S5).

**Figure 2.**
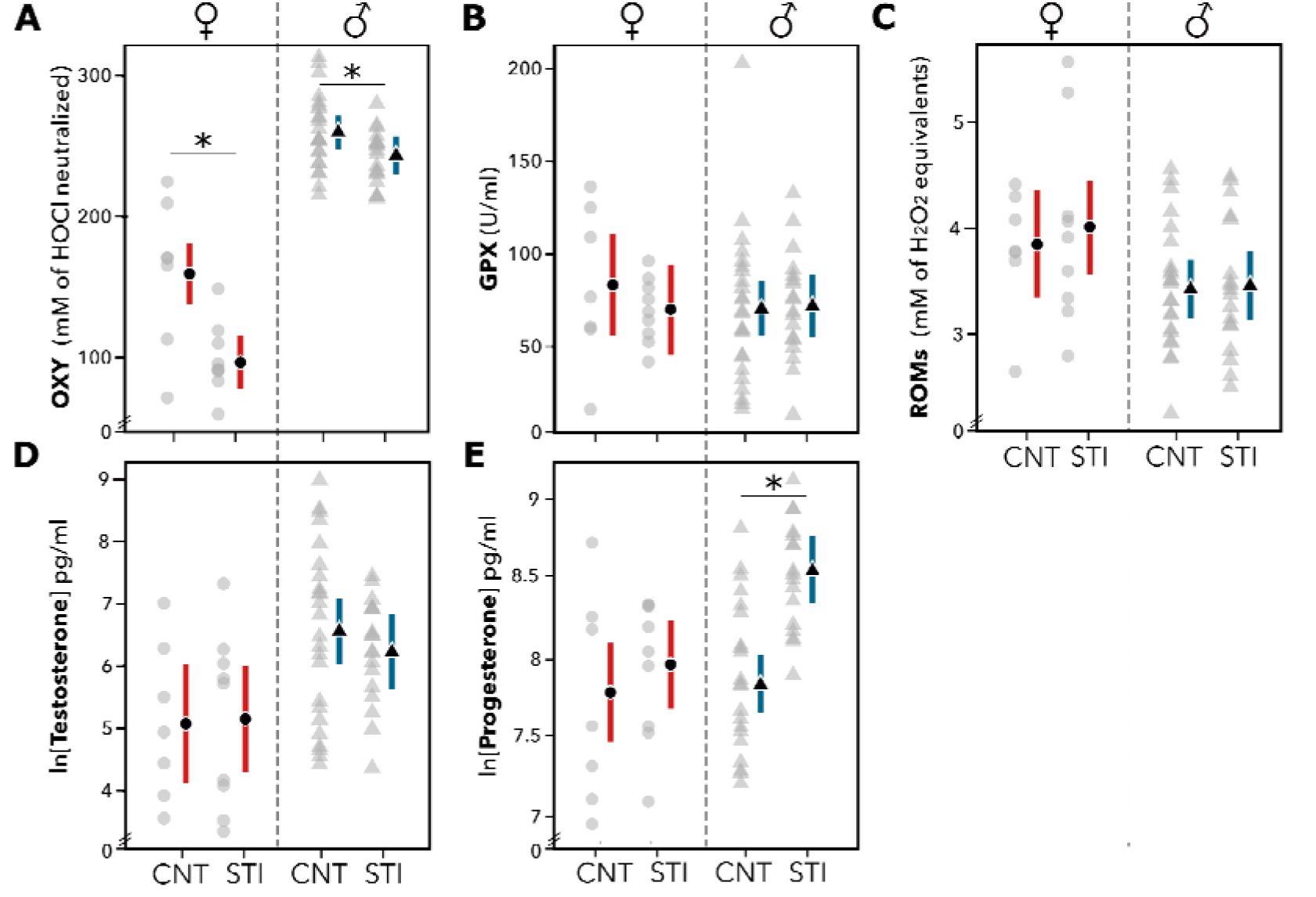
Non-enzymatic antioxidants (OXY), (B) enzymatic antioxidants (GPX), (C) pro-oxidants (ROMs), (D) testosterone and (E) progesterone concentrations measured in control (CNT) and simulated territorial intrusion (STI) female (♀) and male (♂) horneros. Grey circles and triangles show the individual raw data for females and males, respectively. Mean estimates of the model are indicated with black symbols, and 95% credible intervals are represented with colored vertical bars. Statistically meaningful differences are marked with an asterisk.

**Figure 3.**
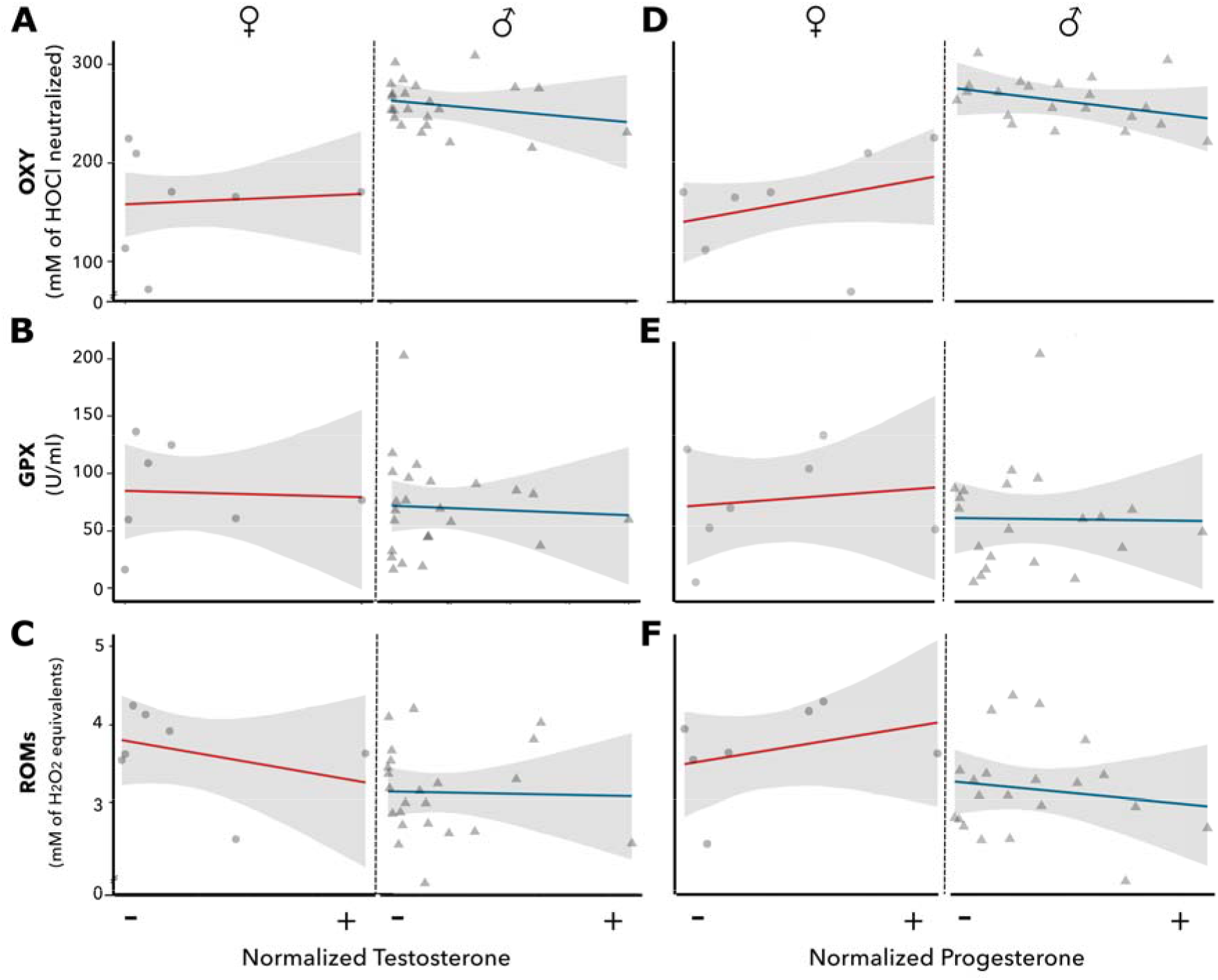
Relationships between the concentrations of non-enzymatic antioxidant (OXY), enzymatic antioxidant (GPX) and pro-oxidants (ROMs) with testosterone (A-C) and progesterone (D-F) in control female (♀) and male (♂) horneros. For each sex, testosterone concentrations were normalized between 0 and 1. Grey circles and triangles show the raw individual data for females and males, respectively. Mean estimates of the model are indicated with colored straight lines, and 95% credible intervals are represented with grey ribbons. There is no relationship between any of the oxidative status markers and testosterone or progesterone concentrations.

### 3.2 Effect of acute conspecific aggressive behavior on the concentrations of oxidative markers, testosterone and progesterone

Both sexes had lower levels of OXY after 20 minutes of simulated territorial intrusion compared to control individuals (Figure 2A; *p(dif.)_females_* > 99.99%; *p(dif.)_males_* > 99.99%). However, the treatment affected females and males differently: the difference in OXY concentrations between control and STI was much stronger in females than in males (females’ OXY differed by 39.62% between treatments, while in males this difference was 6.18%; *p(dif.*) = 99.50%). We found no difference in GPX (*p(dif.)_females_* = 76.62%; *p(dif.)_males_* = 44.85%) or ROMs (*p(dif.)_females_* = 31.35%; *p(dif.)_males_* = 44.14%) concentrations between experimental groups (Figure 2 B-C). In relation to steroid hormones, testosterone concentrations did not differ between control and STI groups (*p(dif.)_females_* = 45.56%; *p(dif.)_males_* = 79.65%; Figure 2 D), whereas, only in males, progesterone concentrations of STI males were higher than that of control males (*p(dif.)_females_* < 60.5%; *p(dif.)_males_* < 99.99%; Figure 2E). For details on model estimates see Table S3.

### 3.3 Relationships between acute conspecific aggressive behavior and the concentrations of oxidative markers, testosterone and progesterone

More aggressive females had lower concentrations of antioxidants than less aggressive females (Figure 4A; posterior probability for the slope being < 0: 99.64%), whereas in males, aggressive behavior and OXY concentrations were unrelated (Figure 4A; posterior probability for the slope being < 0: 52.42%). For both sexes, aggression levels were unrelated to GPX, ROMs, testosterone (Figure 4B) and progesterone (Figure 4C) concentrations (i.e., for each variable, the posterior probability for the slope being < 0 was smaller than 90 %). For details on model estimates see Table S6.

**Figure 4.**
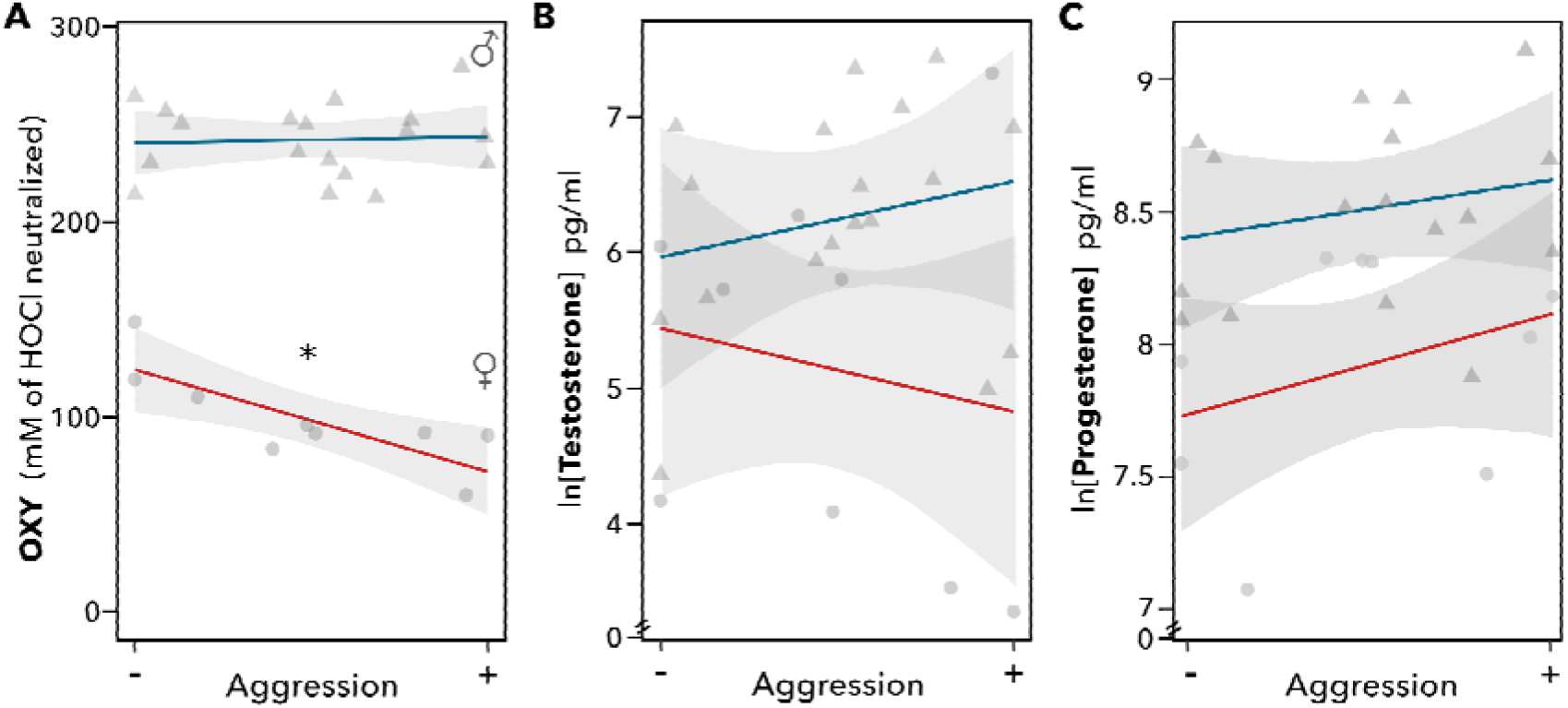
Relationships between the concentrations of (A) non-enzymatic antioxidant (OXY), (B) testosterone and (C) progesterone concentrations with conspecific aggressive behavior in female (♀) and male (♂) horneros. Circles and triangles show the individual raw data for females and males, respectively. Mean estimates of the model are indicated with solid lines (♀: red line; ♂: blue line), and 95% credible intervals are indicated with a shaded area. Statistically meaningful effects are marked with an asterisk.

## 4. Discussion

We experimentally tested if conspecific aggressive behavior induced an acute change in the oxidative state of female and male wild birds, and whether this effect was mediated by testosterone or progesterone. Our results provide the first evidence that an acute increase in conspecific aggressive behavior (i.e., for 20 mins) perturbates the oxidative status of an individual by decreasing the concentration of non-enzymatic antioxidants in plasma. This perturbation in oxidative condition was not related to the concentrations of testosterone or progesterone and was much stronger in females than males. Moreover, lower antioxidant concentrations were correlated to higher aggressive behavior in females, but not in males.

### 4.1. Sex differences in the concentrations of oxidative markers, testosterone and progesterone in control birds

Control females had lower concentrations of non-enzymatic antioxidants (OXY) and a trend for higher concentrations of reactive oxygen species (ROMs) than control males (Figure 2A & 2C). These results were unexpected given that female horneros are less aggressive (Diniz et al. 2018; Mentesana et al., 2020), have lower concentrations of testosterone (Figure 2D) and have similar body condition (Table S2) than males. Also, estrogens, which are generally present at higher concentrations in females close to egg laying compared to males, can increase antioxidant defenses (Halifeoglu et al., 2003). One possible explanation for the sex difference in oxidative status might be that females and males face different reproductive challenges during the early reproductive period (i.e., when females are about to lay eggs). In birds, the resting metabolic rate of females can increase by up to 27% during the period of egg production (Nilsson & Råberg, 2001). Such an increase in metabolic rate could potentially explain the females’ lower levels of OXY and the higher levels of ROMs (*p(dif.*) = 94.75%; Figure 2C) compared to males. The egg laying period is also physiologically demanding since some of the compounds that mothers allocate into their eggs are limited in their availability (Møller et al., 2000; Surai 2002; Hulbert & Abbott, 2011). Antioxidants, such as vitamin E and carotenoids, cannot be synthesized *de novo* (e.g., Surai et al., 2001) and so are of dietary origin. Therefore, another possible and non-exclusive explanation for the difference in the oxidative state between sexes, could be associated with females allocating non-enzymatic antioxidant components into the egg yolk (Royle et al., 1999; Mentesana et al., 2019) to protect the developing embryo (Surai, 2002).

### 4.2. Effect of conspecific aggressive behavior on oxidative status

Because reactive oxygen species are a primary product of aerobic metabolism, it is often assumed that behaviors that increase metabolic rate, such as aggression, produce higher rates of reactive oxygen species (Costantini 2008; Powers & Jackson, 2008; Skrip & McWilliams, 2016; Cooper-Mullin & McWilliams, 2016). However, we found no difference in ROMs between control and STI individuals in either sex (Figure 2C). In contrast, an acute increase in aggressive behavior decreased the antioxidant capacity in male and female horneros (Figure 2A): the concentration of OXY was lower in STI than in control birds. These results have two possible explanations: horneros were able to avoid oxidative damage (as measured by ROMs) by depleting non-enzymatic antioxidants or horneros remobilized non-enzymatic antioxidants among tissues (see below). The concentration of enzymatic antioxidants (GPX) did not differ between experimental groups. GPX concentrations increase with behaviors that associated to increased metabolic rate and are sustained over time (e.g., low-medium endurance training or parental provisioning, Powers and Jackson, 2008; Casagrande & Hau 2018). Such behaviors performed over short-term periods of time can also induce *di novo* protein synthesis at the cellular level; however, protein synthesis occurs in the time scale of hours (e.g., Hernandez et al. 2000). It is thus possible that the upregulation of GPX remained undetectable after 20 minutes of acute aggressive interactions.

Aggressive behaviors can also perturbate the oxidative status indirectly through an increase in steroid hormone concentrations (Schantz et al., 1999; Alonso-Alvarez et al., 2007; Isaksson et al., 2011). However, our results suggest that neither testosterone nor progesterone mediated the change in oxidative condition after an acute increase in conspecific aggressive behaviors. On the one hand, control and STI birds had similar testosterone concentrations; thus, indicating that the difference in oxidative state observed between experimental groups is not mediated by this steroid hormone. In line with our results, the relationship between aggressive phenotypes and oxidative status was also independent of plasma testosterone concentrations in both male and female wild-caught lizards (Isaksson et al., 2011). On the other hand, we found that male, but not female, STI birds had higher progesterone concentrations than controls; yet, our data suggests that aggressive behavior, oxidative status and progesterone are not interrelated for this species: females suffered a much stronger STI-induced decrease in OXY than males (see below), in neither sex there was a relationship between aggressive behavior and progesterone concentrations, and we found no correlation between OXY and progesterone in female and male STI horneros (Pearson’s correlation; R_female_ =-0.48, p-value=0.19; R_male_ = 0.27, p-value=0.29).

The effect that acute conspecific aggressive behaviors had on OXY concentrations was much stronger in females than in males (i.e., STI females had 40% lower OXY concentrations than control females, whereas STI males had only 6% lower OXY concentrations than controls). This sex difference could be given by a differential re-allocation of antioxidants to protect: gametes (Rojas Mora et al., 2017), the developing embryo (see above; Shantz et al., 1999; Surai et al., 2001; Parolini et al. 2017), and/or those tissues with the greatest oxygen consumption (e.g., muscles; Ji 2008). Sex differences could also be explained by other physiological pathways besides an increase in activity levels (see 4.3 for further discussion). The implications of these sex differences in non-enzymatic antioxidant depletion will depend on the life-history stage that the individuals are in (Speakman et al., 2015). Repeated aggressive interactions experienced by females during egg laying may come at a cost to the deposition of substances into the egg that could affect their fitness. For example, an exercised group of female zebra finches (*Taeniopygia guttata*) that had poorer oxidative status than non-exercised birds after flight training, decreased the deposition of antioxidants (i.e., lutein) into the egg yolk (Skrip et al., 2016). However, birds can also recover their circulating non-enzymatic antioxidant capacity by consuming antioxidant-rich foods to avoid negative fitness consequences (e.g., Scrip & McWilliams, 2016). Hence, to understand if aggression-induced perturbations in the oxidative status of an individual have long-lasting functional or fitness consequences, future studies should ideally measure the oxidative condition of an individual not only immediately after facing aggressive interactions but also over repeated interactions, in relation to fitness traits and across different life-history stages.

To the best of our knowledge this is the first study that demonstrates an effect of an acute increase in aggressive behavior (i.e., 20 minutes) on the oxidative status in wild animals. To date, only two other studies investigated the relationship between aggressive behavior and oxidative status by looking at the correlation between the personality trait “aggression” (i.e., over a long-term scale) and the oxidative status of a selected line of mice (Costantini et al., 2008) and white skinks (Isaksson et al., 2011). Interestingly, these two studies support our findings: they also found a relationship between aggression and OXY concentrations, but not with ROMs as previously suggested. Altogether, this suggests that aggression has both short- and long-term consequences on the oxidative status of an individual, and that this occurs through a change in non-enzymatic antioxidants. On a broader scale, these studies show that the relationship between aggression and oxidative status is present across different vertebrate groups (i.e., reptiles, mammals and birds).

### 4.3. Relationships between acute conspecific aggressive behavior and the concentrations of oxidative markers, testosterone and progesterone

Whereas more aggressive females had lower levels of OXY than less aggressive females, no such relationship was found in males. There was also no relationship between aggressive behavior and testosterone or progesterone concentrations in either sex. Together with the previous results, this indicates that mechanisms other than testosterone and progesterone determine the oxidative condition of males and females after conspecific aggressive interactions.

Engaging in aggressive interactions during the nest-building period could also have led to an increase in another steroid hormone: glucocorticoids. An increase in glucocorticoid concentrations can occur through an activation of the hypothalamic-pituitary-adrenal axis to help an individual recover and survive an immediate challenge (Sapolsky et al., 2000). An increase in the concentration of this other steroid hormone after aggressive interactions has been reported both in seasonally territorial (e.g., Charlier et al., 2009) and year-round territorial vertebrates (Landys et al., 2010; Ros et al., 2014). Further, a direct link between glucocorticoids and oxidative state has previously been reported (e.g., Haussmann & Marchetto 2010; Costantini et al., 2011; Haussmannn et al., 2012; Majer et al., 2019), and females seem to be more susceptible to the effects of glucocorticoids on oxidative stress than males (reviewed by Costantini et al., 2011, but see Ouyang et al., 2016). Hence, the interaction between glucocorticoids and the redox system may provide an alternative explanation of our results: aggression may lower OXY concentrations directly because of an increased metabolism, or indirectly via an increase in glucocorticoid concentrations, or both.

## 5. Conclusions

Our study provides the first evidence that an acute increase in aggressive behavior perturbates the oxidative state of female and male wild birds. Our study also suggests that the mechanisms underlying these changes differ between sexes. Until now, the effect of aggression on the oxidative state was investigated in the framework of personality traits (Costantini et al. 2008; Isaksson et al. 2011). Altogether, the existing studies suggest that aggressive behaviors performed over both short and long time scales have direct consequences on the oxidative status of an individual. Remarkably, by generating a change in non-enzymatic antioxidants and not in the concentration of pro-oxidants as frequently suggested. Given that perturbations in the oxidative status can shape central life history traits such as reproduction and longevity, the oxidative state of an individual might be a key physiological mechanism underlying different life-history strategies in individuals that show different levels of aggression.

### Ethical statement

The experimental procedures were approved by the Ethics Committee of Animal Experimentation (CEUA) of the Facultad de Ciencias, Universidad de la República, Uruguay (University of the Republic of Uruguay). Protocol number 186, file 2400-11000090-16.

## Competing interests

We declare no competing interests.

## Funding

We acknowledge funding and support from the International Max Planck Research School (IMPRS) for Organismal Biology, and by Idea Wild that provided field equipment.

## Acknowledgements

We thank Enzo Cavalli and Ernesto Guedes for their assistance in the field, and Monika Trappschuh for conducting the hormone analysis. We also thank the members from the Ethology Lab at the Universidad de la República in Montevideo and especially to Bettina Tassino for their support. Also, we thank all the “INIA Las Brujas” staff for providing us with accommodation and equipment during fieldwork. Furthermore, we are thankful to Juan Carlos Reboreda and the Ecology and Behaviour lab (LEyCA) from the University of Buenos Aires for the logistic support. We also thank Michaela Hau and Manfred Gahr for their valuable support. We thank Laurie O’Neil and Shane McPherson for helping us with the English proof of the manuscript. We are grateful to Stefania Casagrande, Michaela Hau, Wolfgang Goymann and Scott McWilliams for valuable discussions about the study and constructive criticism on previous versions of the manuscript. Lastly, we would like to thank the two anonymous reviewers for constructive comments that helped improving this article.

## Notes

### Competing Interest Statement

The authors have declared no competing interest.

### Summary of Updates

Whole text, and supplementary information was included in the main text

