## Supplementary materials for "Acute aggressive behavior perturbates the oxidative status of a wild bird independently of testosterone and progesterone"

### *1. Supplementary Materials and Methods*

### *Inter-observer reliability test*

Before carrying out the main experiments the two observers performed simulated territorial intrusions (STI) in eight territories that were outside the boundaries of the main study site. Here, instead of registering the activity of the two birds responding to the STI (one bird by each observer), the observers registered the activity of the same bird responding to the STI within each territory. Before initiating the STI both observers would agree on recording the behavior of the bird that would respond first (i.e. First individual approaching the decoy) or the one that would respond second to the STI, and this decision was made at random for each of the eight territories. Also, both observers put special attention on being far enough (>35 m) from each other to prevent hearing what each of them was saying towards the voice recorder. In this way they could avoid any incidental hearing of each other that might influence their observations. After this, we performed statistical models to test if the observers differed in their form of quantifying the birds’ behavior. We performed one model for each trait that was quantified (generalized linear or linear models depending on the nature of the response variable: glm’s for counts and lm’s time variables). The response variable in each model corresponded to 1) response latency, 2) time spent within 5 m of the decoy, 3) time spent on the nest, 4) number of solo songs, 5) number of duets, and 6) number of flights over the decoy. The explanatory variable in each model was the observer (Observer 1/Enzo and Observer 2/Ernesto). See Figure S1 with the raw data, the estimates and the 95% Credible intervals from each model. An absence of statistical difference can be assumed if the estimate for one observer overlaps the 95% CrI of the other. Detailed estimates of the model are in Table S1.

### *Analysis of blood progesterone levels*

Progesterone data for the males were published by Adreani *et al.* (2018). For the females, the same procedure was followed, testosterone was extracted from the plasma and concentrations were determined by radioimmunoassay following Goymann *et al.* (2008). Mean recovery during extraction of the samples were 80 ± 3% (mean ± sd) for females. The detection limit of the hormone assay was 0.39 pg/tube. All samples used in our analyses were above these detection limits. The intra-assay variation, calculated from of an extracted chicken pool, was 12.2%.

### Progesterone statistical analysis

### *Relationship between baseline progesterone and oxidative status*

We ran three linear models to investigate the relationship between baseline oxidative status and progesterone (P4) levels. OXY, GPX, or dROMs were fitted as response variable, and in each model, P4 was fitted as a covariate. Sex and the interaction of sex with P4 levels were included as fixed factors. Since males and females generally differ in their P4 concentrations for each sex we normalized P4 levels between 0 and 1. This allowed us to compare results between sexes.

### *Effect of STI on progesterone levels*

We analyzed the effect that 20 minutes playback had on the progesterone levels of male and female horneros. To do so, we plotted a linear model fitting progesterone (log-transformed) as a response variable. Sex (female vs male), and treatment (CNT vs STI) were included as fixed effects, as well as the interaction between both factors. We also included capture and handling time as covariates to account for the variation that this might add. Preliminary exploration of our data showed that time of the day as well as observer ID did not explain much variance, thus they were not included in the final model.

### *Relationship between progesterone and aggressive behavior*

To study if aggression had an effect on progesterone, we fitted progesterone concentrations as the response variable in a linear model. Progesterone was log-transformed for a better model fit. We used “flights over the decoy” as a covariate in the model since this is the best proxy for aggression during a territorial intrusion in horneros (Adreani *et al.* 2018). In addition, since male and female horneros differ in their aggressive behaviors (i.e., males are more aggressive than females; Diniz *et al.* 2018), for each sex we normalized the behavioral variables between 0 and 1 in order to capture the aggression gradient in each sex and be able to compare the effect between them in one model. Therefore, “normalized flights over the decoy” was included as a covariate in the model. Sex, together with the interaction between aggressive behavior and sex, were fitted as fixed factors. Given that we had a limited sample size and that neither capture nor handling time had an effect on any of the oxidative status parameters or testosterone levels (See Table 1 and Table 2), these variables were not included in the final model.

### *2. Supplementary Figures*

**Fig. S1**. Observer 1 and 2 did not differ in how they scored behaviors. Colored circles show the raw data for each individual bird that was observed. Mean estimates of the model (black dots) and 95% credible intervals (black vertical bars) are also shown.

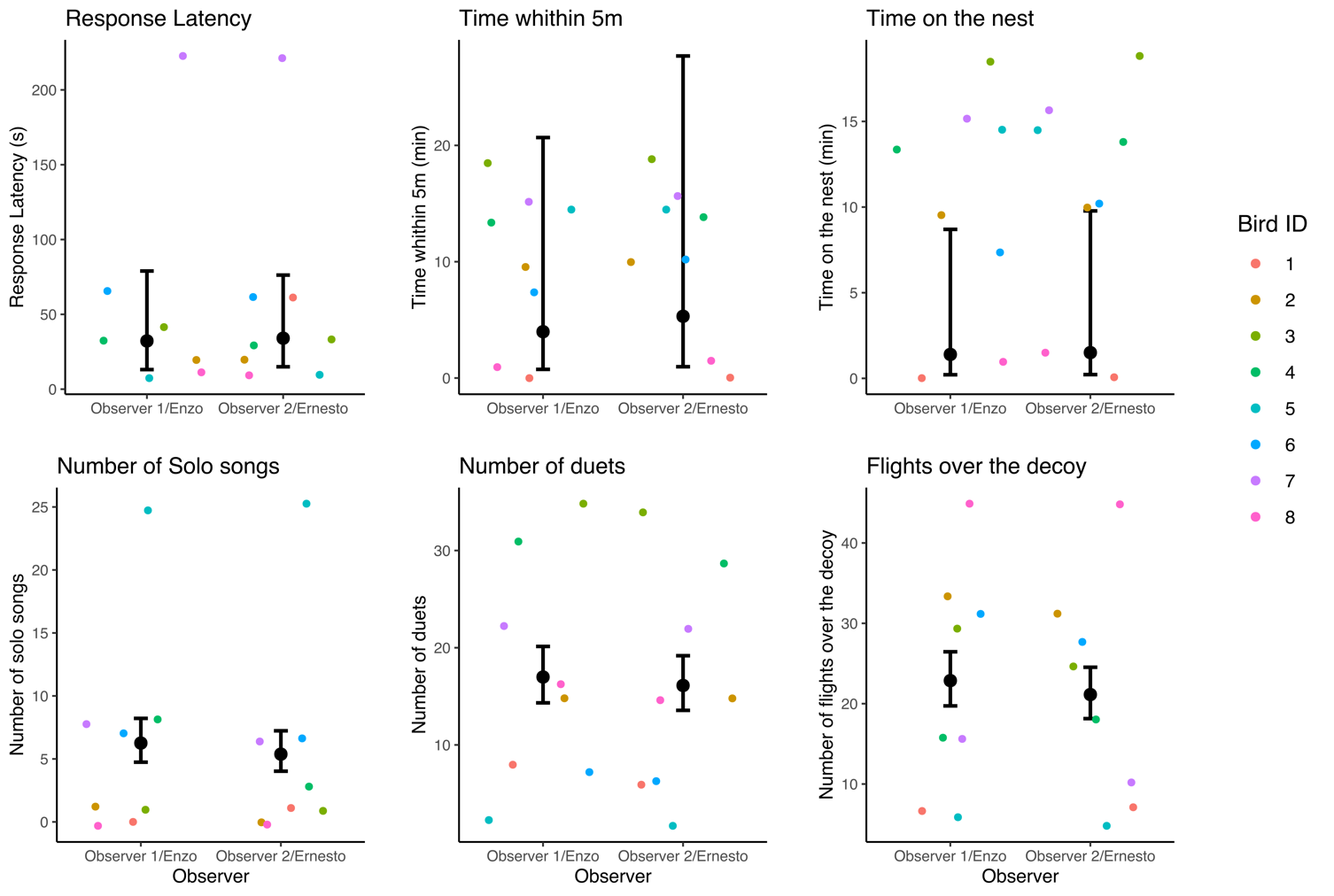

**Fig. S2**. Mean estimates (black shapes) and credible intervals (vertical bars) for the capture times of control (CNT) and STI birds derived from the linear model:

*Capture Time ~ Treatment(CNT vs. STI) * Sex (Female/circles vs. Male/triangles).*

Ten minutes dashed line indicates the minimum time until changes in testosterone (Testo.) can be detected in birds (Wingfield & Wada, 1989). Twenty minutes dashed line indicates the moment in which the playback stopped and the mist nets where unfolded for the STI group; for the CNT birds the mist nets were unfolded since the beginning of the playback and this was stopped immediately after the capture.

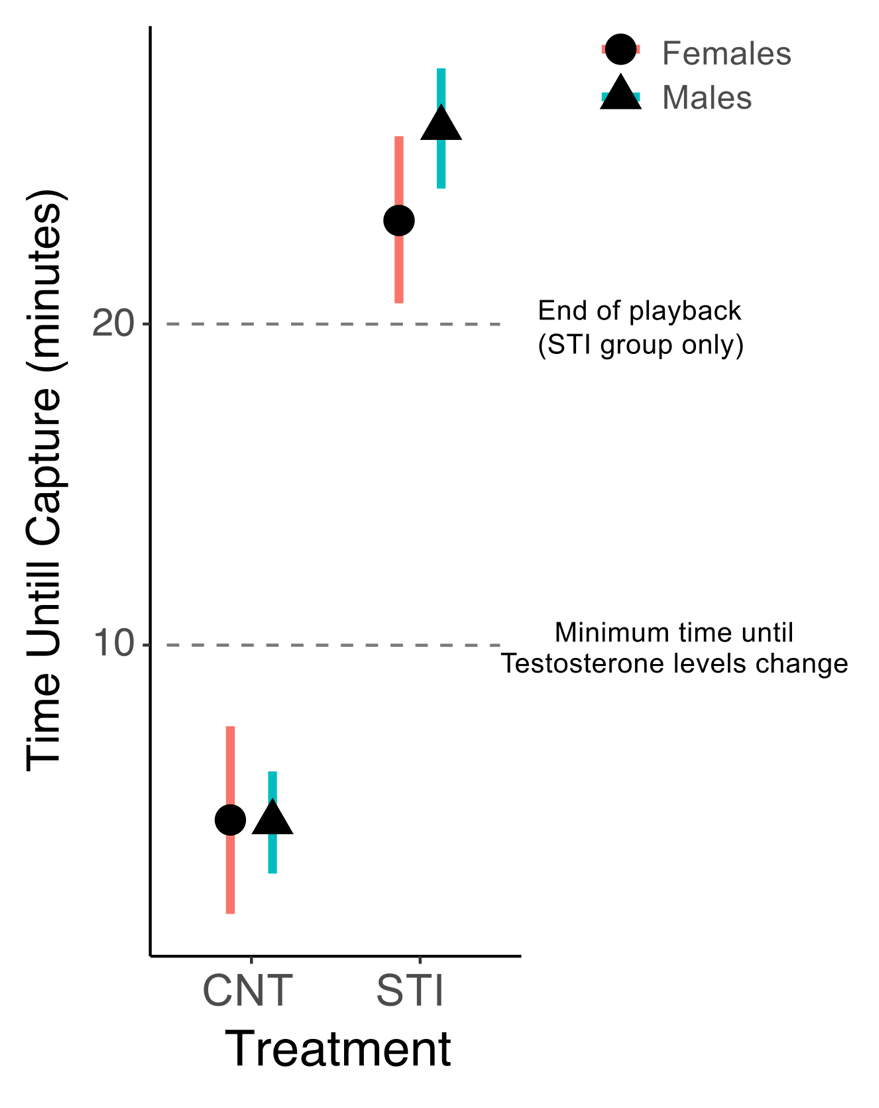

### *3. Supplementary Tables*

**Table S1.** Results from the models comparing the behavior scores of the two observers (observer 1 *vs* 2) showing that observers did not differ in the quantification of the behaviors they measured. Linear mixed effect models were performed for response latency, time within 5 m and time on the nest. Generalized linear mixed models were performed for the count variables: number of solo songs, number of duets and number of flights over the decoy. We present fixed (β) parameters with their 95% credible intervals (CrI) in brackets. A difference between observers can be assumed if zero is not included within the 95% CrI.

|  | Response Latency | Time within 5 m | Time on  Nest | Solo  songs | Duet  songs | Flights over  The decoy |
| --- | --- | --- | --- | --- | --- | --- |
| *Fixed effects β (95% CrI)* | | | | | | |
| Intercept | 3.48  (2.58; 4.34) | 5.47  (3.83; 7.12) | 4.43  (2.57;6.21) | 1.83  (1.55;2.11) | 2.83  (2.66; 3.00) | 3.13  (2.98; 3.27) |
| Observer2 | 0.04  (-1.14; 1.24) | 0.28  (-2.03; 2.62) | 0.10  (-2.40; 2.68) | -0.15  (-0.56;0.25) | -0.05  (-0.29;0.18) | -0.08  (-0.29; 0.12) |

**Table S2.** Results from a linear model estimating fixed effects to explain variation in hornero’s body condition. Tarsus and wing length were only weakly correlated with body mass (*r* < 0.40) and hence we only worked with mass as a proxy of body condition. Treatment (control vs. STI) and sex (female vs. male) were fitted as fixed factors together with the interaction between both factors. We present fixed (β) parameters with their 95% credible intervals (CrI) in brackets. Fixed factors with a statistically meaningful effect can be assumed if zero is not included within the 95% CrI.

|  | Mass |
| --- | --- |
| *Fixed effects β (95% CrI)* | |
| Intercept | 57.77  (54.93; 60.54) |
| Treatment | -0.87  (-4.66; 2.81) |
| Sex | -0.04  (-3.18; 3.19) |
| Treatment * Sex | 0.95  (-3.43; 5.38) |

**Table S3.** Effect of acute aggressive interactions on oxidative stress parameters and testosterone levels (Figure 1). Treatment (control vs. STI), sex (female vs. male) were fitted as fixed factors together with the interaction between both factors. Capture and handling time were included as covariates in the model. We present fixed (β) parameters with their 95% credible intervals (CrI) in brackets. Fixed factors with a statistically meaningful effect (i.e., if zero is not included within the 95% CrI) are presented in bold. Estimates and CrI of ‘0.00’ represent an effect smaller than 0.01.

|  | OXY^1^ | GPX^2^ | dROMs^3^ | Testosterone |
| --- | --- | --- | --- | --- |
| *Fixed effects β (95% CrI)* | | | | |
| Intercept | 173.59  (144.66; 202.07) | 82.55  (46.10; 119.15) | 3.58  (2.90; 4.26) | 5.26  (4.07; 6.49) |
| Treatment | -62.68  (-90.44; -34.99) | -13.27  (-49.40; 23.03) | 0.17  (-0.50; 0.84) | 0.02  (-1.22; 1.24) |
| Sex | 100.23  (76.47; 125.12) | -12.82  (-44.55; 18.75) | -0.42  (-0.98; 0.17) | 1.36  (0.31; 2.42) |
| Treatment * Sex | 46.29  (12.61; 80.05) | 14.80  (-28.34; 57.79) | -0.13  (-0.95; 0.65) | -0.21  (-1.68; 1.21) |
| Capture time^4^ | -1.12  (-3.27; 0.99) | 0.44  (-2.11; 3.01) | 0.00  (-0.04; 0.05) | -0.02  (-0.10; 0.06) |
| Handling time^5^ | -0.03  (-0.09; 0.03) | -0.00  (-0.08; 0.07) | 0.00  (-0.00; 0.00) | -0.00  (-0.00; 0.00) |

^1^ Non-enzymatic antioxidants in plasma.

^2^ Enzymatic antioxidant in red blood cells.

^3^ Reactive oxygen metabolites in plasma.

^4^ Time (seconds) since mist nets were unfolded until an individual was captured.

^5^ Time (seconds) since an individual was trapped into the mist net until blood sample was taken.

**Table S4.** Results from a linear model estimating fixed effects to explain variation in baseline oxidative stress parameters and testosterone levels of control male and female hornero (Figure 2). Sex (female vs. male) was fitted as a fixed factor together with capture and handling time as covariates in the model. We present fixed (β) parameters with their 95% credible intervals (CrI) in brackets. Fixed factors with a statistically meaningful effect (i.e., if zero is not included within the 95% CrI) are presented in bold. Estimates and CrI of ‘0.00’ represent an effect smaller than 0.01.

|  | OXY^1^ | GPX^2^ | dROMs^3^ |
| --- | --- | --- | --- |
| *Fixed effects β (95% CrI)* | | | |
| Intercept | 157.81  (125.19; 191.36) | 85.14  (43.41; 127.15) | 3.96  (3.35; 4.54) |
| Normalized  Testosterone | 10.74  (-67.07; 87.75) | -5.91  (-105.36; 93.23) | -0.57  (-1.92; 0.83) |
| Sex | 105.58  (67.47; 142.89) | 13.69  (-61.62; 33.92) | -0.50  (-1.16; 0.18) |
| Testo*Sex | -32.13  (-127.43; 63.44) | -1.76  (-123.03; 122.03) | 0.50  (-1.22; 2.19) |

^1^ Non-enzymatic antioxidants in plasma.

^2^ Enzymatic antioxidant in red blood cells.

^3^ Reactive oxygen metabolites in plasma

**Table S5.** Effect of aggressive behavior on non-enzymatic antioxidant concentration (OXY) and testosterone of female and male hornero (Figure 3, dROMs and GPX are not shown in the figure)**.** Relationship between aggression and oxidative stress parameters and testosterone levels in horneros. Sex (female vs. male) was fitted as fixed factors. Aggressive behavior (“flights over the decoy”) was included as a covariate. The interaction between both factors was also included in the model. We present fixed (β) parameters with their 95% credible intervals (CrI) in brackets. Fixed factors with a statistically meaningful effect (i.e., if zero is not included within the 95% CrI) are presented in bold.

|  | OXY^1^ | GPX^2^ | dROMs^3^ | Testosterone |
| --- | --- | --- | --- | --- |
| *Fixed effects β (95% CrI)* | | | | |
| Intercept | 123.98  (102.35; 145.64) | 8.19  (6.27; 10.10) | 1.97  (1.75; 2.19) | 5.43  (4.19; 6.67) |
| Aggressive behavior^4^ | -52.14  (-87.87; -16.55) | 0.19  (-3.04; 3.45) | 0.03  (-0.36; 0.39) | -0.61  (-2.67; 1.47) |
| Sex | 116.48  (89.59; 143.49) | -0.53  (-2.97; 1.90) | -0.17  (-0.46; 0.12) | 0.52  (-1.05; 2.14) |
| Aggressive behavior * Sex | 55.32  (11.09; 99.63) | 1.12  (-2.89; 5.13) | 0.07  (-0.40; 0.56) | 1.17  (-1.53; 3.77) |

^1^ Non-enzymatic antioxidants in plasma.

^2^ Enzymatic antioxidant in red blood cells.

^3^ Reactive oxygen metabolites in plasma.

^4^ Number of flights over the decoy per minute was used as a proxy for aggressive behavior (Adreani *et al.*, 2018) and normalized between 0 and 1 within each sex.

**Table S6.** Results from a linear model estimating the effect of baseline progesterone levels on the oxidative stress parameters of rufous hornero. Sex (female vs. male) was fitted as a fixed factor. Progesterone concentration was included as a covariate. The interaction between both factors was also included in the model. We present fixed (β) parameters with their 95% credible intervals (CrI) in brackets. Fixed factors with a statistically meaningful effect can be assumed if zero is not included within the 95% CrI.

|  | OXY^1^ | GPX^2^ | dROMs^3^ |
| --- | --- | --- | --- |
| *Fixed effects β (95% CrI)* | | | |
| Intercept | 140.91  (106.27;174.97) | 77.97  (29.94; 126.59) | 3.64  (2.98; 4.30) |
| Progesterone | 59.00  (-13.95; 130.49) | 15.62  (-86.08; 117.29) | 0.52  (-0.86; 1.93) |
| Sex | 130.15  (89.49; 170.04) | -9.50  (-66.29; 47.53) | -0.15  (-0.91; 0.61) |
| Progesterone * Sex | -91.34  (-181.15; -1.40) | -17.68  (-142.88; 105.47) | -0.83  (-2.52; 0.87) |

^1^ Non-enzymatic antioxidants in plasma.

^2^ Enzymatic antioxidant in red blood cells.

^3^ Reactive oxygen metabolites in plasma

**Table S7.** Results from the linear model estimating the effect of acute aggressive interactions on hornero’s progesterone levels. Progesterone was log-transformed for a better fit. Treatment (control vs. STI), sex (female vs. male) were fitted as fixed factors together with the interaction between both factors. Capture and handling time were included as covariates in the model. We present fixed (β) parameters with their 95% credible intervals (CrI) in brackets. Fixed factors with a statistically meaningful effect (i.e., if zero is not included within the 95% CrI) are presented in bold. Estimates and CrI of ‘0.00’ represent an effect smaller than 0.01.

|  | Progesterone |
| --- | --- |
| *Fixed effects β (95% CrI)* | |
| Intercept | 7.23  (6.80; 7.67) |
| Treatment | 0.17  (-0.26; 0.59) |
| Sex | 0.05  (-0.33; 0.42) |
| Treatment * Sex | 0.55  (0.04; 1.07) |
| Capture time^1^ | 0.00  (-0.02; 0.03) |
| Handling time^2^ | 0.00  (0.00;0.00) |

^1^ Time (seconds) since mist nets were unfolded until an individual was captured.

^2^ Time (seconds) since an individual was trapped into the mist net until blood sample was taken.

**Table S8.** Results from a linear model estimating the effect of aggressive behavior on the progesterone levels (log-transformed) of rufous hornero. Sex (female vs. male) was fitted as a fixed factor. Aggressive behavior (“flights over the decoy”) was included as a covariate. The interaction between both factors was also included in the model. We present fixed (β) parameters with their 95% credible intervals (CrI) in brackets. Fixed factors with a statistically meaningful effect (i.e., if zero is not included within the 95% CrI) are presented in bold.

|  | Progesterone |
| --- | --- |
| *Fixed effects β (95% CrI)* | |
| Intercept | 7.73  (7.28; 8.18) |
| Aggressive behavior^1^ | 0.38  (-0.35; 1.12) |
| Sex | **0.67**  **(0.12; 1.225)** |
| Aggressive behavior * Sex | -0.16  (-1.08; 0.77) |

^1^ Number of flights over the decoy per minute was used as a proxy for aggressive behavior (Adreani *et al.*, 2018) and normalized between 0 and 1 within each sex.

Neotropical ovenbird: Sexual and seasonal variation and adaptive signaling

functions. *Journal of Avian Biology*, **49**, jav‐01637.

Goymann, W., Wittenzellner, A., Schwabl, I. & Makomba, M. (2008) Progesterone modulates aggression in sex-role reversed female African black coucals. *Proceedings of the Royal Society of London B: Biological Sciences*, **275**.

Wingfield, J. C., & Wada, M. (1989). Changes in plasma levels of testosterone during male-male interactions in the song sparrow, Melospiza melodia: time course and specificity of response. Journal of Comparative Physiology A, **166**(2), 189-194.
